# *Mycobacterium abscessus* biofilms have viscoelastic properties which may contribute to their recalcitrance in chronic pulmonary infections

**DOI:** 10.1101/2020.10.20.347252

**Authors:** Erin S. Gloag, Daniel J. Wozniak, Paul Stoodley, Luanne Hall-Stoodley

## Abstract

*Mycobacterium abscessus* is emerging as a cause of recalcitrant chronic pulmonary infections, particularly in people with cystic fibrosis (CF). Biofilm formation has been implicated in the pathology of this organism, however the role of biofilm formation in infection is unclear. Two colony-variants of *M. abscessus* are routinely isolated from CF samples, smooth (*Ma*^Sm^) and rough (*Ma*^Rg^). These two variants display distinct colony morphologies due to the presence (*Ma*^Sm^) or absence (*Ma*^Rg^) of cell wall glycopeptidolipids (GPLs). We hypothesized that *Ma*^Sm^ and *Ma*^Rg^ variant biofilms might have different biophysical and mechanical properties, including stiffness, viscosity and elasticity. To test this hypothesis, we performed uniaxial mechanical indentation, and shear rheometry on *Ma*^Sm^ and *Ma*^Rg^ colony-biofilms. We identified that *Ma*^Rg^ biofilms were significantly stiffer than *Ma*^Sm^ under a normal force, while *Ma*^Sm^ biofilms were more pliant compared to *Ma*^Rg^, under both normal and shear forces. Furthermore, using theoretical indices of mucociliary and cough clearence, we identified that *M. abscessus* biofilms may be more resistant to mechanical forms of clearance from the lung, compared to other common pulmonary pathogens, such as *P. aeruginosa.* Thus, the mechanical properties of *M. abscessus* biofilms may contribute to the persistent nature of pulmonary infections caused by this organism.

## Introduction

Nontuberculous mycobacteria (NTM) are ubiquitously found in soil and water and are difficult to eradicate ^1–3^. NTM lung infections are an emerging health threat in the general population, outpacing tuberculosis in the US and Europe ^4–8^. NTM have grown to be a common complication in cystic fibrosis (CF), where NTM now rank as the third most frequent cause of lung infection in people with CF ^9,10^. *Mycobacterium abscessus*, a NTM that presents as extracellular bacterial biofilm aggregates in the lung ^11,12^, has emerged as an important pathogen in people with bronchiectasis and chronic lung diseases ^3,13–15^. *M. abscessus* causes intractable infections and considerable morbidity and mortality in individuals suffering from chronic obstructive pulmonary disease (COPD), primary ciliary dyskinesia (PCD) and CF ^8,16,17^.

*M. abscessus* infection is particularly challenging in people with CF ^5,7,9,10,17^. CF is caused by mutations in the cystic fibrosis transmembrane conductance regulator (*cftr*) gene, resulting in a spectrum of effects on CFTR function and subsequent disease. The precise mechanisms of how defective CFTR leads to recurrent infection is the topic of some debate ^18^. However, it is widely understood that the accumulation of viscous mucus in the airway obstructs the effective clearance of bacteria, leading to bacterial growth and recurrent progressive infections, inflammation, bronchiectasis and, eventually, respiratory failure.

Furthermore, people with CF infected with *M. abscessus* are younger and have a poorer prognosis than those infected with other NTM, leading to severe bronchiectasis and increased mortality ^9,10^. Antibiotic therapy for *M. abscessus* can be as prolonged as for multi-drug resistant tuberculosis, lasting 12 months or longer. Over 60% of CF patients infected with *M. abscessus* have adverse effects or toxicities that limit antibiotic therapy ^7,13,14,16^. The mechanisms underlying the rising incidence of *M. abscessus* infection in people with CF however, remain ill-defined ^8,19^.

*M. abscessus* has two distinct colony variants, based on the presence (smooth morphotype; *Ma*^Sm^) or absence (rough morphotype; *Ma*^Rg^) of cell wall glycopeptidolipids (GPL) ^20,21^. The surface GPL phenotypes influence important aspects of *M. abscessus* pathobiology, including biofilm formation, aggregation, host cell invasion, macrophage bactericidal activity, and virulence ^20,22^. However, the *Ma*^Rg^ variant is considered more virulent than the *Ma*^Sm^ because it is associated with rapid deterioration of lung function and progression of lung disease ^10^.

A prevailing view of chronic *M. abscessus* infection is that *Ma*^Sm^ is a noninvasive, biofilm-forming, persistent phenotype, and *Ma*^Rg^ is an invasive phenotype that is unable to form biofilms. We have previously shown, however, that *Ma*^Rg^ is hyper-aggregative and is capable of forming biofilm-like aggregates, which are significantly more tolerant than planktonic *M. abscessus* to acidic pH, hydrogen peroxide (H_2_O_2_), or antibiotic treatment ^20^. These studies indicate that development of biofilm-like aggregates and biofilms contribute to the persistence of *M. abscessus* in the face of antimicrobial agents, regardless of morphotype. Antibiotic regimens that reliably cure *M. abscessus* infections are lacking. A better understanding of this organism and its ability to cause persistent pulmonary infections is needed to explore new treatment strategies ^23,24^.

The study of biofilm mechanics has gained interest, as it is being realized that these properties are important for the understanding of biofilm biology and how biofilms respond to chemical and mechanical forms of treatment, including removal ^25^. Furthermore, changes in colony morphology, not only has been correlated to differences in biofilm formation ^26^, but also differences in biofilm mechanics ^27,28^. We therefore hypothesized that the biofilms of *Ma*^Sm^ and *Ma*^Rg^ might have different biophysical and mechanical properties, due to their distinct colony morphologies and GPL composition, and that these properties may help explain putative differences in persistence and virulence in these two phenotypes. In the present study, we used mechanical indentation and shear rheometry to evaluate viscoelastic properties of *Ma*^Sm^ and *Ma*^Rg^ biofilms. Mechanical indentation analysis indicated that *Ma*^Rg^ biofilms were significantly stiffer than *Ma*^Sm^ biofilms. Shear rheometry analysis further indicated that *Ma*^Sm^ biofilms were significantly more pliant than *Ma*^Rg^ biofilms. However, dynamic oscillatory frequency sweep analysis showed that, in both variants, the viscoelastic response was dominated by elastic behavior over viscous behavior, across the tested range. Notably, the biofilms of each variant had theoretical clearance indices ^29^ that predict mucociliary or cough clearance mechanisms may be ineffective in facilitating the mechanical removal of *Ma*^Sm^ and *Ma*^Rg^ biofilms from the lung. These data support the hypothesis that biofilm viscoelasticity could contribute to the virulence and persistence of biofilm-associated infefctions ^25,30^, and provides novel insight into why *M. abscessus* may be such a persistent pathogen in airway infection in people with CF.

## Results

### *Ma* colony-biofilms maintain the distinct morphologies of *Ma*^Sm^ and *Ma*^Rg^ variants

*M. abscessus* is a rapidly growing NTM, typically showing colony morphology by 4 days on agar media. To test the hypothesis that biofilms of *Ma*^Sm^ and *Ma*^Rg^ variants might differ in their mechanical properties, uniaxial indentation and shear rheology was performed on 4 day *M. abscessus* colony-biofilms. As this is the first time that we have used this biofilm model for *M. abscessus,* we examined the colony-biofilms after 4 days of growth (Fig 1). Macroscopically, biofilm formation was evident, covering the filter. Biofilm morphology maintained the characteristic phenotypes of each variant ^20^. That is, biofilms of *Ma*^Sm^, had a smooth, almost mucoid apparence (Fig 1A), while *Ma^Rg^* biofilms showed a cauliflower-like morphology (Fig 1B) in agreement with our previous study showing cording at the periphery of *Ma^Rg^* colonies ^20^. Importantly, when these colony-biofilms were disrupted, and plated to observe single cells, the number of cells within the biofilm was similar across the two variants, and no reversion across phenotypes was observed during this time (Fig 1C), demonstrating that during the 4 day biofilm growth period, each variant was stable.

**Figure 1:**
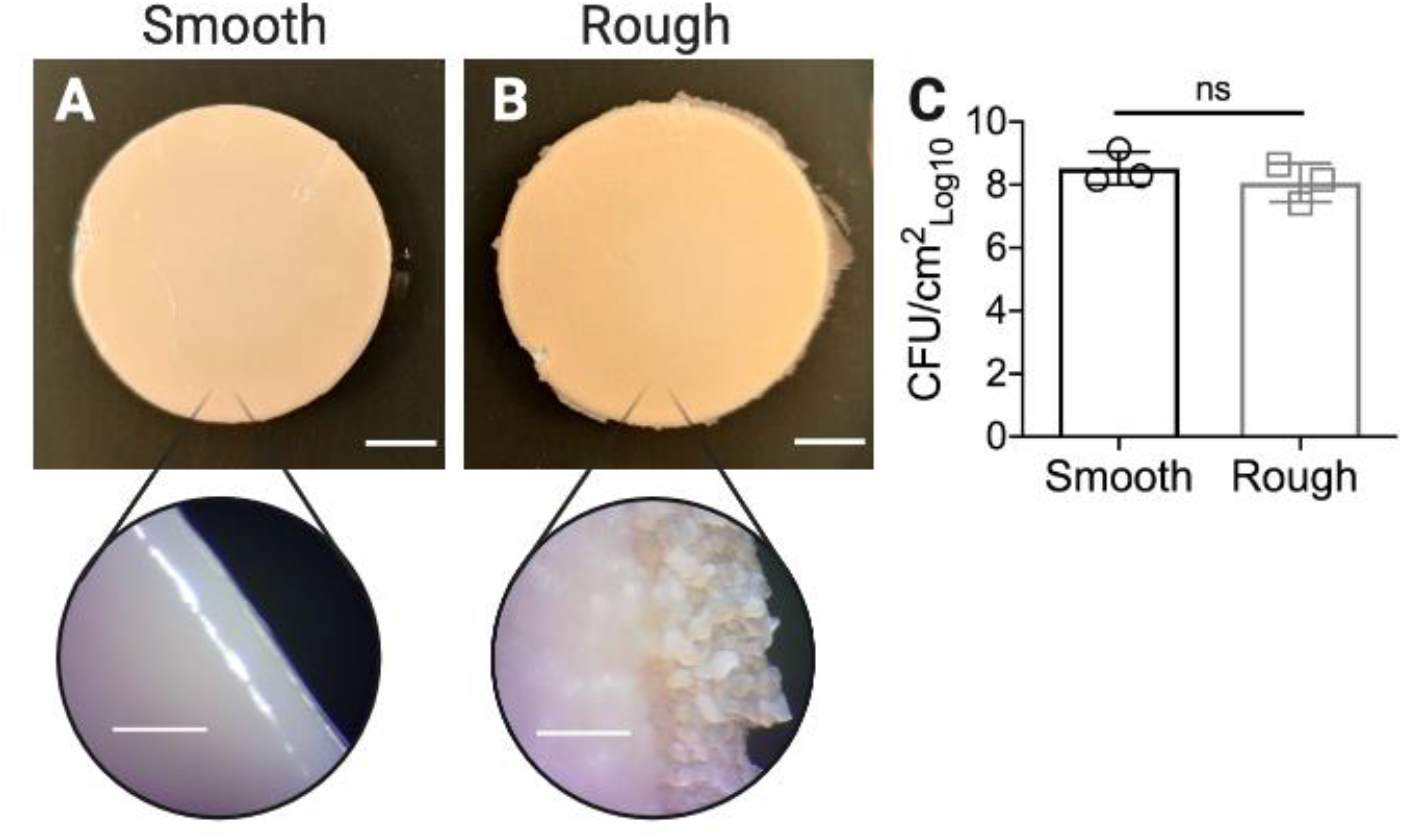
Morphology of *Ma^Sm^* and *Ma^Rg^* colony-biofilms. **(A)** *Ma^Sm^* and **(B)***Ma^Rg^* were grown on nitrocellulose membranes for 4 days, transferring the membranes onto fresh media after 48h. Colony-biofilms were then imaged to visualize the macroscopic morphology. Scale bar indicates 5mm, and 0.5mm for the zoomed insets. **(C)**To assess the biomass and if each variant was stable during biofilm formation, 4 day *M. abscessus* biofilms were enumerated for CFUs. Evidence of phenotypic changes of the colony morphologies for either was not observed at 37ºC over 4 days. Statistical analysis was performed using a Student’s t-test; ns indicated not significant. N=3.

### *Ma*^Rg^ biofilms are stiffer than *Ma*^Sm^ biofilms in uniaxial compression

To determine the stiffness of *M. abscessus* biofilms under a normal force, uniaxial indentation was performed on 4 day *M. abscessus* colony-biofilms. During this analysis, biofilms are compressed and the force required is measured. This analysis is also used to determine the biofilm thickness, which revealed that biofilms of each variant were of similar thickness (Fig 2A), consistent with the observation that biofilms had a similar number of CFUs (Fig 1C). Stress-strain curves revealed that *Ma*^Sm^ and *Ma*^Rg^ biofilms displayed a ‘J-shaped’ curve, where the biofilms became progressively stiffer as they were compressed (Fig 2B). Stress-strain curves also revealed that there were mechanical differences between *Ma*^Sm^ and *Ma*^Rg^ biofilms (Fig 2B). To quantify these differences, the Young’s modulus was determined from the lower linear portions of the curve, which revealed that the Young’s modulus of *Ma*^Rg^ biofilms was approximately two-fold greater than *Ma*^Sm^ biofilms. This indicates that *Ma*^Rg^ biofilms were significantly stiffer under uniaxial compression, compared to *Ma*^Sm^ biofilms (p<0.0001) (Fig 2C).

**Figure 2:**
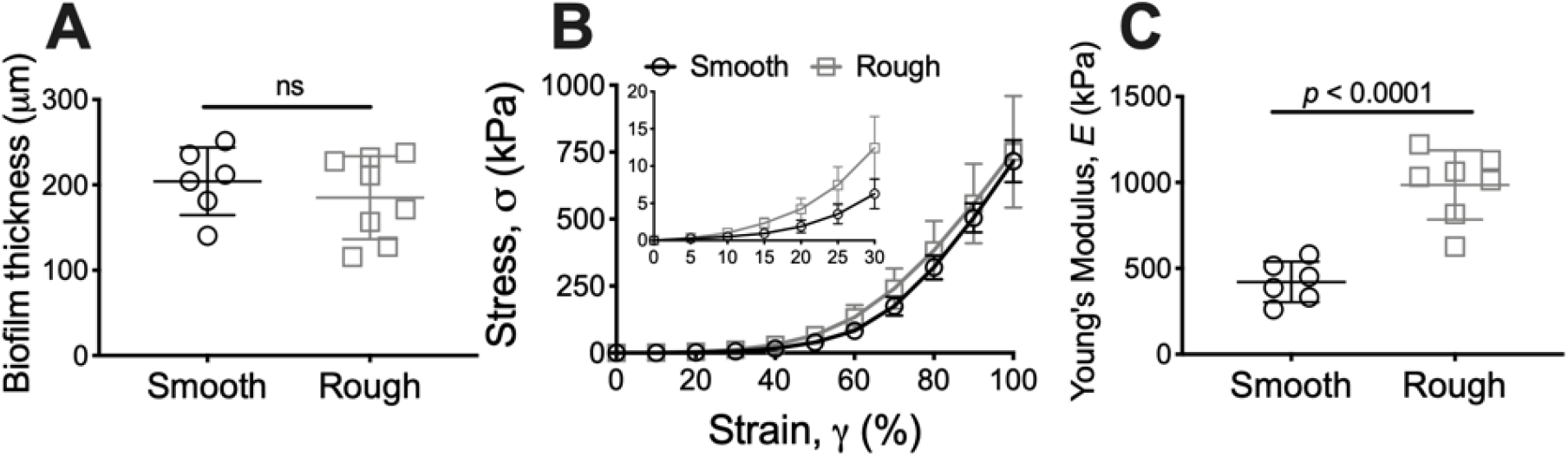
*Ma*^Rg^ colony-biofilms are stiffer compared to *Ma*^Sm^ biofilms in compression. **(A)** Thickness and **(B)**stress-strain curves of 4 day *M. abscessus* colony-biofilms determined from uniaxial indentation analysis. Inset in **(B)** depicts 0-30% strain. **(C)**Young’s modulus of *M. abscessus* biofilms, determined from the lower linear potion of the force-displacement curve, corresponding to 0-30% strain. N=4; data presented as individual data points with mean ± SD. Statistical analysis was performed using a Student’s t-test; ns indicates not significant.

### *M. abscessus* biofilms are highly elastic and *Ma*^Sm^ biofilms are more pliant than *Ma*^Rg^ under shear

To examine the mechanical properties of *M. abscessus* colony-biofilms in further detail, *Ma*^Sm^ and *Ma*^Rg^ biofilms were analyzed using spinning disc rheology. Oscillatory strain sweeps were performed, where the oscillatory strain was incrementally increased and the storage and loss moduli measured, which reflect the elastic and viscous response, respectively ^25^. The measured storage (G’) and loss (G”) moduli plateaus were similar for *Ma*^Sm^ and *Ma*^Rg^ biofilms (Fig 3A, B), suggesting that biofilms of each variant behaved similarly within the linear viscoelastic region. From this analysis we determined the yield strain, which represents the strain where the biofilm integrity begins to break down, due to the increasing applied strain, and transitions to fluid-like behavior characteristic of a viscoelastic fluid ^31^. Interestingly, the yield strain of *Ma*^Sm^ biofilms was significantly greater than *Ma*^Rg^ biofilms (p=0.0098) (Fig 3C). This suggests that *Ma*^Sm^ biofilms are more pliant than *Ma*^Rg^ biofilms, and that *Ma*^Sm^ biofilms can be deformed to a greater extent before cohesive failure occurs. This is also consistent with the Young’s modulus of these biofilms, which indicated that *Ma*^Sm^ biofilms can be compressed more readily compared to *Ma*^Rg^ biofilms (Fig 2C).

**Figure 3:**
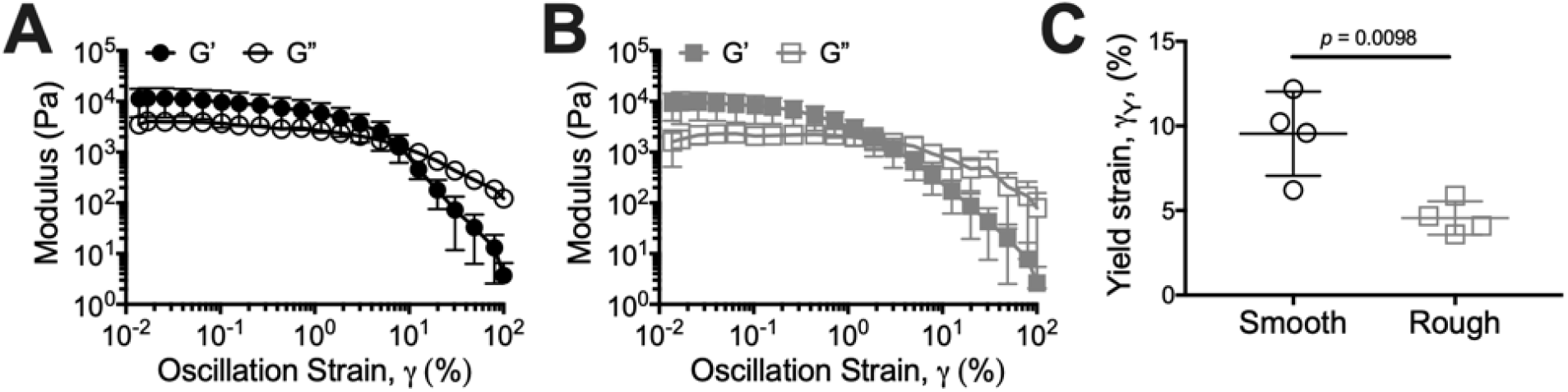
*Ma*^Sm^ colony-biofilms are more pliant than *Ma*^Rg^ biofilms in shear. Strain sweep profiles of 4 day **(A)** *Ma*^Sm^ and **(B)** *Ma*^Rg^ colony-biofilms. N=4; date presented as mean ± SD. **(C)** Yield strain of *M. abscessus* colony-biofilms determined from **(A)** and **(B)**. Data presented as individual data points with mean ± SD. Statistical analysis was performed using a Student’s t-test.

Oscillatory frequency sweeps were also performed, where the oscillatory frequency was incremented under a consistent applied strain, and the storage and loss moduli measured. Across the range of frequencies analyzed, the storage and loss moduli were relatively independent of frequency, with the storage modulus greater than the loss modulus for both *M. abscessus* variant biofilms (Fig 4A, B). This indicates that both *Ma*^Sm^ and *Ma*^Rg^ biofilms displayed elastic behavior that was dominant across the analyzed conditions. There were no significant differences between the mechanical behavior of *Ma*^Sm^ and *Ma*^Rg^ biofilms using this analysis (Fig 4C, D). However, compared to *Ma*^Sm^ biofilms, *Ma*^Rg^ biofilms trended toward a reduced loss modulus across the analyzed frequencies (Fig 4D).

**Figure 4:**
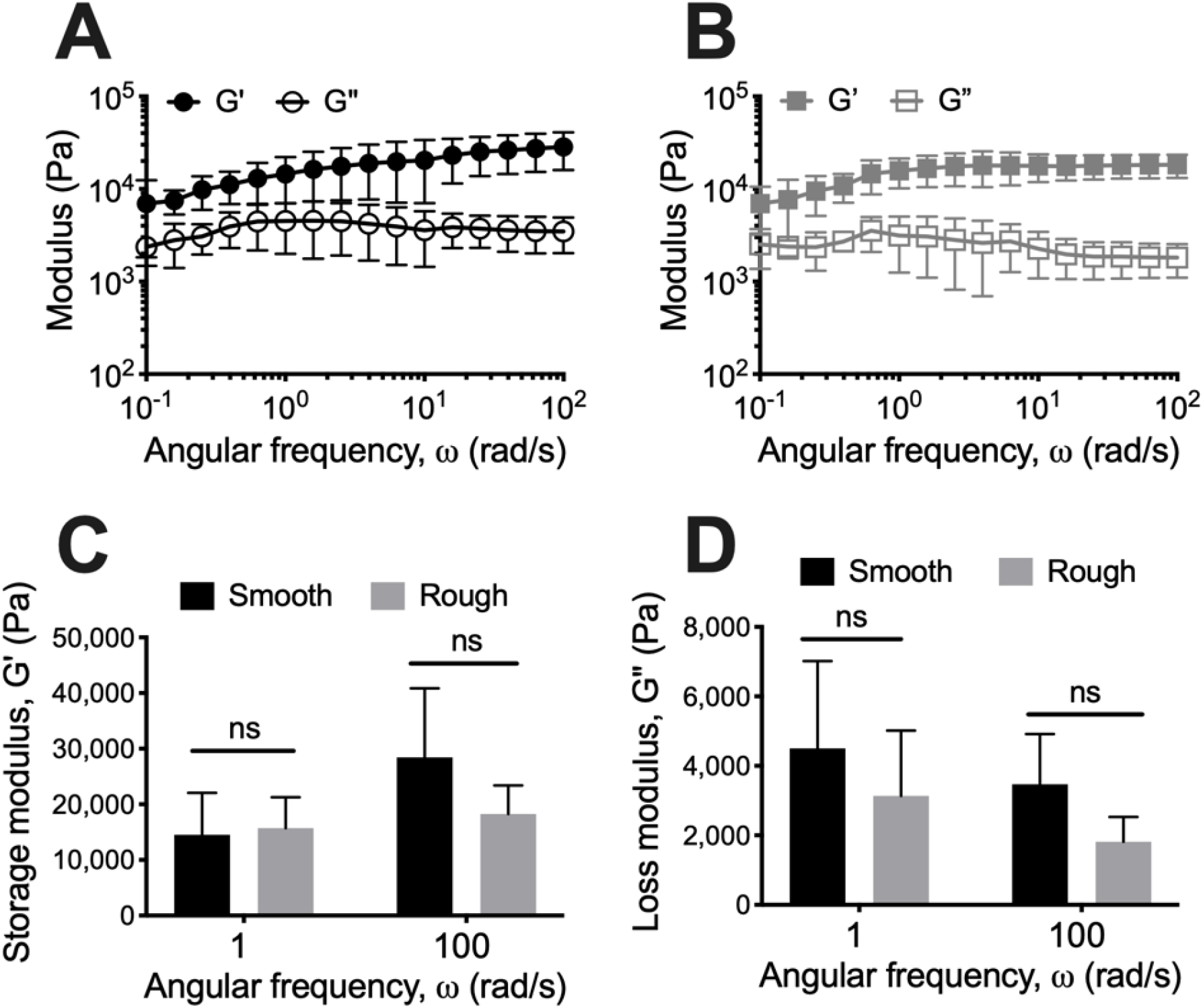
*M. abscessus* colony-biofilms are highly elastic. Frequency sweep profiles of 4 day **(A)** *Ma*^Sm^ and **(B)** *Ma*^Rg^ colony-biofilms. **(C)** Storage and **(D)** and loss moduli values at 1 and 100 rad/s. N=4; data presented as mean ± SD. Statistical analysis was performed using a Student’s t-test; ns indicates not signficant.

### Theoretical indices suggest that *M. abscessus* biofilms, independent of morphotype, may resist clearance from the lung

To determine if the mechanical properties of *M. abscessus* biofilms may correlate with recalcitrance of *M. abscessus* in pulmonary infections, we calculated the mucociliary (MCI) and cough (CCI) clearance index, according to equations 2 and 3, using values from the frequency sweep analysis of these biofilms (Fig 4). The MCI and CCI were developed to correlate sputum viscoelasticity to predicted levels of clearance from the lung via either mechanism ^32,33^. The MCI and CCI of *Ma*^Sm^ and *Ma*^Rg^ biofilms were similar. However, due to the highly viscoelastic behavior of both colony variant biofilms (Fig 4A, B), the indices were less than the MCI and CCI reported for sputum collected from people with CF (Fig 5; line).

**Figure 5:**
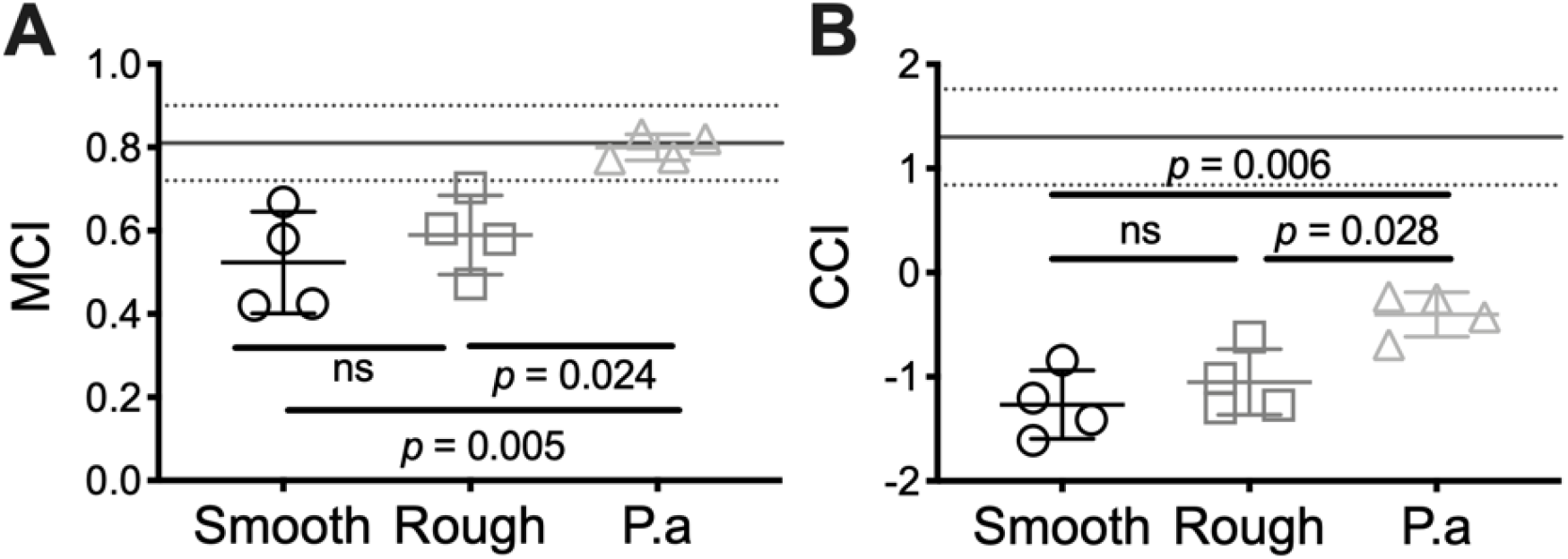
*M. abscessus* colony-biofilms have reduced theoretical mucociliary and cough clearance index. **(A)** Mucociliary and **(B)** cough clearance index of 4-day *M. abscessus* and *P. aeruginosa* (P.a) colony-biofilms. Lines at **(A)** and **(B)** are the MCI (0.81 ±0.09) and CCI (1.3 ± 0.46), respectively, of sputum collected from patients with CF, previously determined by ^29^. N=4; Data presented as individual data points with mean ± SD. Statistical analysis was performed using a one-way ANOVA with a Tukey’s post-hoc test; ns indicates not significant.

To compare *M. abscessus* biofilms to the biofilms of another common pathogen causing chronic pulmonary infections in people with CF, we analyzed 4 day wild type *Pseudomonas aeruginosa* colony-biofilms ^34^. *P. aeruginosa* is considered a model organism for biofilm formation, and we have previously determined the MCI and CCI for this organism ^27^. Using the same testing conditions for *M. abscessus* biofilms described here, we performed oscillatory frequency sweeps on 4 day wild type *P. aeruginosa* colony-biofilms (See Supplmentary Results; Fig S1A), and similarly determined the MCI and CCI for comparison (Fig 5). The indices of *Ma*^Sm^ and *Ma*^Rg^ biofilms were significantly less than that determined for wild type *P. aeruginosa* biofilms (Fig 5). This suggests that *M. abscessus* biofilms may be more resistant to mechanical clearance from the lung, compared to other common pulmonary pathogens, such as *P. aeruginosa*.

## Discussion

Our findings presented here provide novel insight into potential reasons why *M. abscessus* is a persistent and difficult to treat pulmonary pathogen. We also provide the first description of the mechanical properties of *M. abscessus* biofilms, comparing the biofilms of smooth and rough morphology variants that are commonly isolated from patients with pulmonary infection. We found that *Ma*^Rg^ colony-biofilms were stiffer under normal forces (Fig 2C), while *Ma*^Sm^ colony-biofilms were more pliable under both normal and shear forces (Fig 2C, 3C). In constrast to the uniaxial indentation analysis (Fig 2), oscillatory frequency sweeps indicated that there were no significant difference in the viscoelasticity of *Ma^Sm^* and *Ma^Rg^* biofilms under shear forces (Fig 4). This indicates that *M. abscessus* biofilms were anisotropic, in that they have different mechanical properties when exposed to either a normal or shear force. Viscoelasticity is a property of biofilms that allows dissipation of the force, rather than breaking or cohesive failure, when exposed to mechanical forces such as shear. The elastic property allows a biofilm to return to its shape during intermittent perturbations, whereas the viscous component enables biofilms to flow similar to liquids when forces are sustained ^25,35^. These properties along with the extracellular polymeric substance (EPS) matrix contribute to biofilm stability.

Earlier research by others suggested that only *Ma*^Sm^ variants formed biofilms and speculated that GPL expression enhanced sliding motility in CF mucus, making it better adapted to behave as a colonizing, biofilm-forming phenotype ^36,37^. However, we have previously shown that the rough morphotype is hyper-aggregative ^20^, indicating that each colony morphology variant is capable of forming biofilm-like aggregates. The finding here that *Ma*^Sm^ colony-biofilms were more pliable under both normal and shear forces (Fig 2C, 3C) suggests that GPL likely affects the mechanical properties of *M. abscessus* biofilms. Further studies are needed to better understand the relationship of *M. abscessus* cell wall determinants, such as lipids, GPLs, and EPS with bioflm function.

Uniaxial indentation, revealed that *Ma*^Sm^ and *Ma*^Rg^ biofilms each displayed a ‘J-shaped’curve in response to increasing compression (Fig 2B). This response is typical of biological materials, and has previously been observed for a number of different types of bacterial biofilms ^27,38–40^. The Young’s modulus of *Ma*^Sm^ and *Ma*^Rg^ colony-biofilms, determined here, is similar to that reported for *Streptococcus mutans* hydrated biofilms ^41^, *P. aeruginosa* colony-biofilms ^27^, and surface adhered *Staphylococcus aureus, Staphylococcus epidermidis* and *Streptococcus salivarius* cells^42^.

Interestingly, *Ma*^Sm^ and *Ma*^Rg^ colony-biofilms have a storage modulus approximately 10-fold greater than that previously observed for other bacterial biofilms (Fig S1A) ^27,28,31,43–51^. This suggests that *M. abscessus* biofilms, regardless of the morphotype, are highly viscoelastic, and much stiffer than biofilms formed by other bacterial pathogens. It would, therefore, be of interest to compare the mechanical properties of other NTM biofilms, as well as other biofilms that have a lipid dominant EPS, to determine the contributions of lipid content to the EPS and to the biofilm mechanical properties.

MCI and CCI are theoretical indices that were developed from *in vitro* lung clearance models, to correlate the viscoelastic properties of expectorated mucus, to predicted levels of clearance from the lung ^32,33^. We previously determined the MCI and CCI of *P. aeruginosa* colony-biofilms, to determine if the viscoelastic properties of bacterial biofilms could impact mechanical clearance from the lung ^27^. We identified that elastic *P. aeruginosa* biofilms had a reduced CCI, while mucoid *P. aeruginosa* biofilms that had a low viscosity, had both a reduced MCI and CCI ^27^. Here, we determined the MCI and CCI of *M. abscessus* biofilms, and similarly observed that these stiffer biofilms had a negative CCI, indicating that they may resist clearance from the lung by cough (Fig 5B). These findings provide novel insight into potential reasons why *M. abscessus* is a persistent and difficult to treat pulmonary pathogen.

A longitudinal rheological study of sputum from CF patients indicated that both viscosity and elasticity increase during pulmonary exacerbations, suggesting that CF sputum viscoelasticity is linked to disease state and airflow obstruction ^52^. Our findings provide a novel way to understand *M. abscessus* biofilms. More broadly, the impact of evaluating bacterial biofilm mechanical properties offers new avenues for the treatment of intractable pulmonary infections. Disruption of *M. abscessus* aggregates increased susceptibility to several antibiotics ^23^. Moreover, liposomes, lipid nanoparticles and nanoparticle lipid carriers are being evaluated for pulmonary use ^53^. This approach has been used on biofilms of *Burkholderia cepacia* complex, as well as *S. aureus* and *P. aeruginosa* biofilms *in vitro* ^54,55^. These products not only have the potential to reduce exposure to high concentrations of antimicrobial agents following aerosol administration by reducing toxicity, but particularly relevant to NTM antibiotic therapy, might ameliorate the severe side effects often associated with antimycobacterial treatment. This strategy could further be used to target NTM, which have a high lipid content, to facilitate cohesive failure of biofilm aggregates in the lung, in adddtion to delivering targeted antibiotics.

## Materials and Methods

### *M. abscessus* culture and biofilm model

Colony-biofilms were grown as previously described with modifications ^27^. Briefly, *M. abscessus* cultures were prepared by inoculating a 10μL loop of isolated *Ma*^Sm^ or *Ma*^Rg^ from 7H10 agar plates into 7H9 without tween media and a single cell suspension prepared as described ^20^. Cultures were normalized to an OD_600_ of 0.15–0.2 and 100μL was pipetted onto sterile nitrocellulose filter membranes (25mm, 0.45μm pore size; Millipore) and incubated at 37C, 5% CO_2_ under humidified conditions for 4 days. Colony-biofilms of PAO1 *P. aeruginosa* were grown as previously described^27^.

Colony-biofilms were imaged using a Stereo Microscope (AmScope) fitted with a Microscope Digital Color CMOS camera (AmScope). Images were processed in FIJI ^56^. Figure was complied in Biorender.com.

To assess the number of CFU/cm^2^ on sterile membranes after 4 days, colony-biofilms were scraped and transferred to 5mL of 7H9+tween 80 and vortexed with glass beads to disperse cells from the biofilm ^20^. Cell suspensions were serially diluted and enumerated for CFUs in triplicate on 7H10 agar plates. To dertermine if either variant converted/reverted to another colony variant during biofilm growth, colony morphology was also assessed and verified by sub-culturing three colonies per replicate to ensure that the colony morphology was stable. CFUs were plated to 10^8^ dilution and colonies were streaked to isolation to validate colony variants.

### Rheometry apparatus

A Discovery Hybrid Rheometer-2 with a heat exchanger attached to the Peltier plate (TA Instruments) was used for all rheological measurements. For uniaxial indentation and spinning disc measurements, the rheometer was fitted with an 8mm and 25mm sand-blasted Smart Swap geometry, respectively. All measurements were performed at 37°C, with the Peltier plate covered with a moist Kimwipe to prevent dehydration of the colony-biofilm. TRIOS v4 (TA instruments) software was used.

### Uniaxial indentation measurements

Uniaxial indentation measurements were performed under compression using an approach rate of 1μm/s. Two colony-biofilms were analyzed per biological replicate (total of 4 biofilms analyzed), with two measurements per biofilm. The point of contact with the biofilm was determined to be where the force began to increase after the initial point of pull-on adhesion. Force-displacement curves were converted to stress-strain curves, for ease of comparison, as previously described^27^. The Young’s modulus was calculated according to equation 1 ^57^:

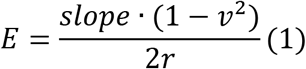

where *v* is the assumed Poisson’s ratio of a biofilm (*v* = 0.5) ^39^ and *r* is the radius of the geometry. The slope is of the force-displacement curve, and was taken at the region corresponding to 0-30% strain.

### Spinning disc rheology

To normalize for differences in biofilm thickness across replicates, prior to analysis, colony-biofilms were compressed to a normal force of 0.01N. For all analyses a total of 4 biofilms were analyzed, 2 per biological replicate.

Strain sweeps were performed by incrementing the oscillatory strain from 0.01-100% at a frequency of 1Hz. The yield strain was taken where the storage (G’) and loss (G”) modulus intersected. Frequency sweeps were performed by incrementing the oscillating frequency from 0.1-100rad/s at a constant strain of 0.1%. This strain was determined to be within the linear viscoelastic region of the strain sweeps for both *Ma*^Sm^ and *Ma*^Rg^ colony-biofilms.

Theoretical mucociliary clearance index (MCI) and cough clearance index (CCI) were calculated according to equations 2 and 3, respectively, using the relationship between the complex modulus (G*) and tanδ values determined from frequency sweep measurements ^3, 2,33,58^. The MCI and CCI was calculated from values determined at an angular frequency of 1rad/s and 100rad/s, respectively.

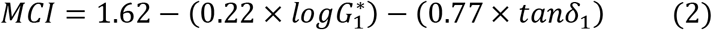

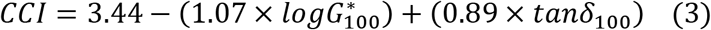

### Statistical analysis

Data are presented as mean ± SD. Statistical significance was determined using either an One-way ANOVA with a Tukey’s post-hoc test or a Student’s t-test. Analyses were performed using GraphPad Prism v.7 (Graphpad Software). Statistical significance was determined using a p-value <0.05.

## Data Availability

All data generated or analysed during this study are included in this published article (and its Supplementary Information files).

## Author contributions

ESG and LHS performed the experiments. ESG, LHS and PS performed the data analysis and interpretation. ESG, DJW, PS and LHS wrote the manuscript.

## Acknoweldgements

Funding to LHS was provided by the Ohio State University College of Medicine Office of Research Bridge Funding Program and by the Cystic Fibrosis Foundation (HALLST18I0). ESG was funded by an American Heart Association Career Development Award (19CDA34630005). DJW and PS were funded by the National Institute of Health (R01AI134895 and R01AI143916; DJW) (R01GM124436; PS).

## References

1 Wallace Jr, R. J., Brown, B. A. & Griffith, D. E. Nosocomial outbreaks/pseudo outbreaks caused by nontuberculous mycobacteria. Annual review of microbiology 52, 453–490 (1998).

2 Larsen, M. H. et al. The Many Hosts of Mycobacteria 8 (MHM8): A conference report. Tuberculosis 121, 101914 (2020).

3 Griffith, D. E. et al. An official ATS/IDSA statement: diagnosis, treatment, and prevention of nontuberculous mycobacterial diseases. American journal of respiratory and critical care medicine 175, 367–416 (2007).

4 Salsgiver, E. L. et al. Changing epidemiology of the respiratory bacteriology of patients with cystic fibrosis. Chest 149, 390–400 (2016).

5 Floto, R. A. et al. Audit, research and guideline update: US Cystic Fibrosis Foundation and European Cystic Fibrosis Society consensus recommendations for the management of non-tuberculous mycobacteria in individuals with cystic fibrosis: executive summary. Thorax 71, 88 (2016).

6 Fleshner, M. et al. Mortality among patients with pulmonary non-tuberculous mycobacteria disease. The International Journal of Tuberculosis and Lung Disease 20, 582–587 (2016).

7 Swenson, C., Zerbe, C. S. & Fennelly, K. Host variability in NTM disease: implications for research needs. Frontiers in microbiology 9, 2901 (2018).

8 Wu, M.-L., Aziz, D. B., Dartois, V. & Dick, T. NTM drug discovery: status, gaps and the way forward. Drug discovery today 23, 1502–1519 (2018).

9 Park, I. K. & Olivier, K. N. in Seminars in respiratory and critical care medicine. 217 (NIH Public Access).

10 Furukawa, B. S. & Flume, P. A. in Seminars in respiratory and critical care medicine. 383–391 (Thieme Medical Publishers).

11 Fennelly, K. P. et al. Biofilm formation by Mycobacterium abscessus in a lung cavity. American journal of respiratory and critical care medicine 193, 692–693 (2016).

12 Qvist, T. et al. Chronic pulmonary disease with Mycobacterium abscessus complex is a biofilm infection. European Respiratory Journal 46, 1823–1826 (2015).

13 Medjahed, H., Gaillard, J.-L. & Reyrat, J.-M. Mycobacterium abscessus: a new player in the mycobacterial field. Trends in microbiology 18, 117–123 (2010).

14 Griffith, D. E. Mycobacterium abscessus subsp abscessus lung disease:‘trouble ahead, trouble behind…’. F1000prime reports 6(2014).

15 Nessar, R., Cambau, E., Reyrat, J. M., Murray, A. & Gicquel, B. Mycobacterium abscessus: a new antibiotic nightmare. Journal of antimicrobial chemotherapy 67, 810–818 (2012).

16 Wu, U.-I. et al. in Open Forum Infectious Diseases. ofz484 (Oxford University Press US).

17 Daniel-Wayman, S. et al. Advancing translational science for pulmonary nontuberculous mycobacterial infections. a road map for research. American journal of respiratory and critical care medicine 199, 947–951 (2019).

18 Malhotra, S., Hayes, D. & Wozniak, D. J. Cystic fibrosis and Pseudomonas aeruginosa: the host-microbe interface. Clinical microbiology reviews 32(2019).

19 Scott, J. P., Ji, Y., Kannan, M. & Wylam, M. E. Inhaled granulocyte–macrophage colony-stimulating factor for Mycobacterium abscessus in cystic fibrosis. European Respiratory Journal 51(2018).

20 Clary, G. et al. Mycobacterium abscessus smooth and rough morphotypes form antimicrobial-tolerant biofilm phenotypes but are killed by acetic acid. Antimicrobial agents and chemotherapy 62, e01782–01717 (2018).

21 Rhoades, E. R. et al. Mycobacterium abscessus glycopeptidolipids mask underlying cell wall phosphatidyl-myo-inositol mannosides blocking induction of human macrophage TNF-α by preventing interaction with TLR2. The Journal of Immunology 183, 1997–2007 (2009).

22 Pawlik, A. et al. Identification and characterization of the genetic changes responsible for the characteristic smooth-to-rough morphotype alterations of clinically persistent M ycobacterium abscessus. Molecular microbiology 90, 612–629 (2013).

23 Kolpen, M. et al. Biofilms of Mycobacterium abscessus complex can be sensitized to antibiotics by disaggregation and oxygenation. Antimicrobial agents and chemotherapy 64(2020).

24 Yam, Y.-K., Alvarez, N., Go, M.-L. & Dick, T. Extreme Drug Tolerance of Mycobacterium abscessus “Persisters”. Frontiers in microbiology 11, 359 (2020).

25 Gloag, E. S., Fabbri, S., Wozniak, D. J. & Stoodley, P. Biofilm mechanics: implications in infection and survival. Biofilm, 100017, doi.org/10.1016/j.bioflm.2019.100017 (2019).

26 Branda, S. S., Vik, A., Friedman, L. & Kolter, R. Biofilms: the matrix revisited. Trends Microbiol. 13, 20–26 (2005).

27 Gloag, E. S., German, G. K., Stoodley, P. & Wozniak, D. J. Viscoelastic properties of Pseudomonas aeruginosa variant biofilms. Scientific reports 8, 9691, 10.1038/s41598-018-28009-5 (2018).

28 Kovach, K. et al. Evolutionary adaptations of biofilms infecting cystic fibrosis lungs promote mechanical toughness by adjusting polysaccharide production. npj Biofilms and Microbiomes 3, 1 (2017).

29 Dasgupta, B. & King, M. Reduction in viscoelasticity in cystic fibrosis sputum in vitro using combined treatment with nacystelyn and rhDNase. Pediatric pulmonology 22, 161–166, 10.1002/(sici)1099-0496(199609)22:3 (1996).

30 Peterson, B. W. et al. Viscoelasticity of biofilms and their recalcitrance to mechanical and chemical challenges. FEMS microbiology reviews 39, 234–245 (2015).

31 Klapper, I., Rupp, C. J., Cargo, R., Purvedorj, B. & Stoodley, P. Viscoelastic fluid description of bacterial biofilm material properties. Biotechnology and bioengineering 80, 289–296, 10.1002/bit.10376 (2002).

32 King, M. Relationship between mucus viscoelasticity and ciliary transport in guaran gel/frog palate model system. Biorheology 17, 249 (1980).

33 King, M., Brock, G. & Lundell, C. Clearance of mucus by simulated cough. Journal of Applied Physiology 58, 1776–1782 (1985).

34 Ciofu, O., Tolker-Nielsen, T., Jensen, P. O., Wang, H. & Hoiby, N. Antimicrobial resistance, respiratory tract infections and role of biofilms in lung infections in cystic fibrosis patients. Advanced drug delivery reviews 85, 7–23, 10.1016/j.addr.2014.11.017 (2015).

35 Koo, H., Allan, R. N., Howlin, R. P., Stoodley, P. & Hall-Stoodley, L. Targeting microbial biofilms: current and prospective therapeutic strategies. Nature reviews. Microbiology, 10.1038/nrmicro.2017.99 (2017).

36 Howard, S. T. et al. Spontaneous reversion of Mycobacterium abscessus from a smooth to a rough morphotype is associated with reduced expression of glycopeptidolipid and reacquisition of an invasive phenotype. Microbiology (Reading, England) 152, 1581–1590 (2006).

37 Nessar, R., Reyrat, J.-M., Davidson, L. B. & Byrd, T. F. Deletion of the mmpL4b gene in the Mycobacterium abscessus glycopeptidolipid biosynthetic pathway results in loss of surface colonization capability, but enhanced ability to replicate in human macrophages and stimulate their innate immune response. Microbiology (Reading, England) 157, 1187–1195 (2011).

38 Devaraj, A. et al. The extracellular DNA lattice of bacterial biofilms is structurally related to Holliday junction recombination intermediates. Proceedings of the National Academy of Sciences 116, 25068–25077 (2019).

39 Rmaile, A. et al. Microbial tribology and disruption of dental plaque bacterial biofilms. Wear 306, 276–284 (2013).

40 Stoodley, P., Lewandowski, Z., Boyle, J. D. & Lappin-Scott, H. M. Structural deformation of bacterial biofilms caused by short-term fluctuations in fluid shear: an in situ investigation of biofilm rheology. Biotechnology and bioengineering 65, 83–92 (1999).

41 Fabbri, S. et al. Streptococcus mutans biofilm transient viscoelastic fluid behaviour during high-velocity microsprays. Journal of the Mechanical Behavior of Biomedical Materials 59, 197–206 (2016).

42 Chen, Y., Norde, W., van der Mei, H. C. & Busscher, H. J. Bacterial cell surface deformation under external loading. mBio 3, e00378–00312 (2012).

43 Grumbein, S., Opitz, M. & Lieleg, O. Selected metal ions protect Bacillus subtilis biofilms from erosion. Metallomics 6, 1441–1450 (2014).

44 Lieleg, O., Caldara, M., Baumgärtel, R. & Ribbeck, K. Mechanical robustness of Pseudomonas aeruginosa biofilms. Soft matter 7, 3307–3314 (2011).

45 Pavlovsky, L., Younger, J. G. & Solomon, M. J. *In situ* rheology of *Staphylococcus epidermidis* bacterial biofilms. Soft matter 9, 122–131, 10.1039/c2sm27005f (2013).

46 Wloka, M., Rehage, H., Flemming, H.-C. & Wingender, J. Structure and rheological behaviour of the extracellular polymeric substance network of mucoid Pseudomonas aeruginosa biofilms. Biofilms 2, 275–283 (2005).

47 Yan, J. et al. Bacterial Biofilm Material Properties Enable Removal and Transfer by Capillary Peeling. Advanced Materials, 1804153 (2018).

48 Waters, M. S., Kundu, S., Lin, N. J. & Lin-Gibson, S. Microstructure and mechanical properties of in situ Streptococcus mutans biofilms. ACS applied materials & interfaces 6, 327–332, 10.1021/am404344h (2014).

49 Cense, A. W. et al. Mechanical properties and failure of Streptococcus mutans biofilms, studied using a microindentation device. Journal of microbiological methods 67, 463–472, 10.1016/j.mimet.2006.04.023 (2006).

50 Houari, A. et al. Rheology of biofilms formed at the surface of NF membranes in a drinking water production unit. Biofouling 24, 235–240, 10.1080/08927010802023764 (2008).

51 Di Stefano, A. et al. Viscoelastic properties of Staphylococcus aureus and Staphylococcus epidermidis mono-microbial biofilms. Microbial biotechnology 2, 634–641 (2009).

52 Ma, J. T., Tang, C., Kang, L., Voynow, J. A. & Rubin, B. K. Cystic fibrosis sputum rheology correlates with both acute and longitudinal changes in lung function. Chest 154, 370–377 (2018).

53 Cipolla, D., Shekunov, B., Blanchard, J. & Hickey, A. Lipid-based carriers for pulmonary products: preclinical development and case studies in humans. Advanced drug delivery reviews 75, 53–80 (2014).

54 Messiaen, A.-S., Forier, K., Nelis, H., Braeckmans, K. & Coenye, T. Transport of nanoparticles and tobramycin-loaded liposomes in Burkholderia cepacia complex biofilms. PloS one 8, e79220 (2013).

55 Dong, D. et al. Distribution and inhibition of liposomes on Staphylococcus aureus and Pseudomonas aeruginosa biofilm. PloS one 10, e0131806 (2015).

56 Schindelin, J. et al. Fiji: an open-source platform for biological-image analysis. Nature methods 9, 676 (2012).

57 Timoshenko, S. & Goodier, J. Theory of Elasticity third edn, (McGraw Hill Higher Education, 1970).

58 Zayas, J. G., Man, G. C. & King, M. Tracheal mucus rheology in patients undergoing diagnostic bronchoscopy. The American review of respiratory disease 141, 1107–1113 (1990).

