## Supplemental Information for "*Mycobacterium abscessus* biofilms have viscoelastic properties which may contribute to their recalcitrance in chronic pulmonary infections"

Erin S. Gloag[1], Daniel J. Wozniak[1,2], Paul Stoodley[1,3,4], Luanne Hall-Stoodley[1*]

### Supplemental Results

<u>Rheology testing parameters can influence the measured mechanical behavior of bacterial biofilms</u>

Here, we also wanted to compare the mechanical properties of *M. abscessus* biofilms to that of another common pulmonary pathogen. We have previously performed an extensive analysis on the mechanical properties of *Pseudomonas aeruginosa* biofilms [1], which is also an an important pathogen causing pulmonary infections in people with cystic fibrosis [2]. We therefore we analyzed 4 day wild type *P. aeruginosa* colony-biofilms using the same rheological parameters used to analyze the *M. abscessus* biofilms. Interestingly, under these conditions (37°C and constant strain of 0.1%), *P. aeruginosa* biofilms displayed increased elastic behavior, compared to what we had previously measured (25°C and constant stress of 0.5Pa [approximately 2% strain] [1]). As such the mucociliary (MCI) and cough (CCI) clearance index for *P. aeruginosa* colony-biofilms determined here, were lower than what we had previously reported [1]. We therefore repeated the frequency sweep analysis, using the testing parameters of our previous study. Under these conditions the viscoelastic properties and MCI and CCI of *P. aeruginosa* colony-biofilms were similar to those we had previously observed [1] (Fig S1B, C), further illustrating how the dynamic nature of biofilm mechanics and how the testing conditions can influence the measured mechanical behaviour.

### Supplemental Figure Legend

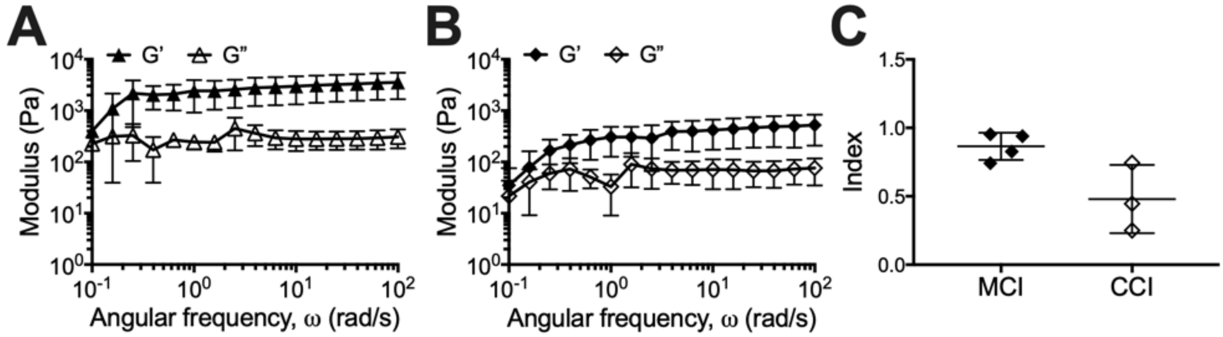


**Supplemental Figure 1: Analysis of *P. aeruginosa* colony-biofilms.** Frequency profiles of wild type *P. aeruginosa* colony-biofilms measured at (A) 37°C and strain of 0.1%, as per the analysis of *M. abscessus* biofilms and at (B) 25°C and stress of 0.5Pa (equivalent to 1.8 ± 0.4% strain) as per our previous analysis [1]. (C) Mucociliary and cough clearance index of P. aeruginosa colony-biofilms determined from the frequency sweep data in (B).

### Supplemental References

1 Gloag, E. S., German, G. K., Stoodley, P. & Wozniak, D. J. Viscoelastic properties of Pseudomonas aeruginosa variant biofilms. *Scientific reports* **8**, 9691, doi:10.1038/s41598-018-28009-5 (2018).

2 Ciofu, O., Tolker-Nielsen, T., Jensen, P. O., Wang, H. & Hoiby, N. Antimicrobial resistance, respiratory tract infections and role of biofilms in lung infections in cystic fibrosis patients. *Advanced drug delivery reviews* **85**, 7-23, doi:10.1016/j.addr.2014.11.017 (2015).
